# Frequency-Blended Diffusion Models for Synthetic Generation of Biologically Realistic Splice Site Sequences

**DOI:** 10.1101/2025.06.16.660047

**Authors:** Espoir Kabanga, Seonil Jee, Arnout Van Messem, Wesley De Neve

## Abstract

We present a frequency-blended diffusion framework for generating biologically realistic splice site sequences. Our approach combines a U-Net-based denoising diffusion probabilistic model with conditional nucleotide frequency priors derived from real donor (5’) and acceptor (3’) splice site sequences in *Arabidopsis thaliana* and *Homo sapiens*. By guiding the generative process with position-specific empirical base frequencies, the model captures both local sequence motifs and long-range dependencies that are critical for realistic splice site representation. We evaluate the synthetic sequences through direct assessments (sequence logos, GC content, nucleotide conservation) and indirect functional tests using state-of-the-art splice site classifiers (SpliceRover, SpliceFinder, DeepSplicer, Spliceator). Our results show that frequency blending substantially improves motif conservation, compositional fidelity, and model transferability. This work establishes frequency-blended diffusion as a promising strategy for generating high-quality nucleotide sequences for modeling, benchmarking, and data augmentation in genomics research.

## I. Introduction

DNA splicing is a critical eukaryotic process in which introns are removed from pre-mRNA and exons are joined to form mature mRNA [1]. Accurate identification of donor (5’) and acceptor (3’) splice sites is essential for understanding gene structure. Donor sites typically feature a `GT`dinucleotide at the exon-intron boundary, and acceptor sites typically display an `AG`motif at the intron-exon boundary [2], both surrounded by conserved sequence patterns [3].

Generating synthetic DNA sequences with authentic splice site characteristics can help augment datasets, test predictive algorithms, and enable comparative genomics [4]. Traditional approaches, such as position weight matrices (PWMs) and Markov models, capture local nucleotide dependencies but assume statistical independence across positions or are restricted to learning short-range dependencies [5], [6]. These limitations hinder their ability to model the complex global sequence patterns found in biological sequences. To address such challenges, recent work has explored incorporating experimental feedback to guide biological sequence generation and improve functional outcomes [7]. Moreover, recent deep generative models, particularly diffusion models such as DNA-Diffusion [8] and DiscDiff [9], offer a powerful framework to learn both local motifs and long-range dependencies.

However, standard diffusion models do not incorporate the positional constraints characteristic of splice sites. We introduce a conditional diffusion framework that blends nucleotide frequency information with the generative process of diffusion models by incorporating position-specific base compositions from real alignments. This strategy guides the model with empirical priors while leveraging the diffusion process to capture higher-order dependencies. Our contributions are: (1) a novel conditional diffusion model for splice site sequence generation, (2) its application to generate realistic splice site sequences in (*Arabidopsis thaliana* and *Homo sapiens*), and (3) improved biological fidelity over unconditional diffusion models. Figure 1 illustrates our pipeline: data preprocessing, diffusion training, and sequence generation.

**Fig. 1.**
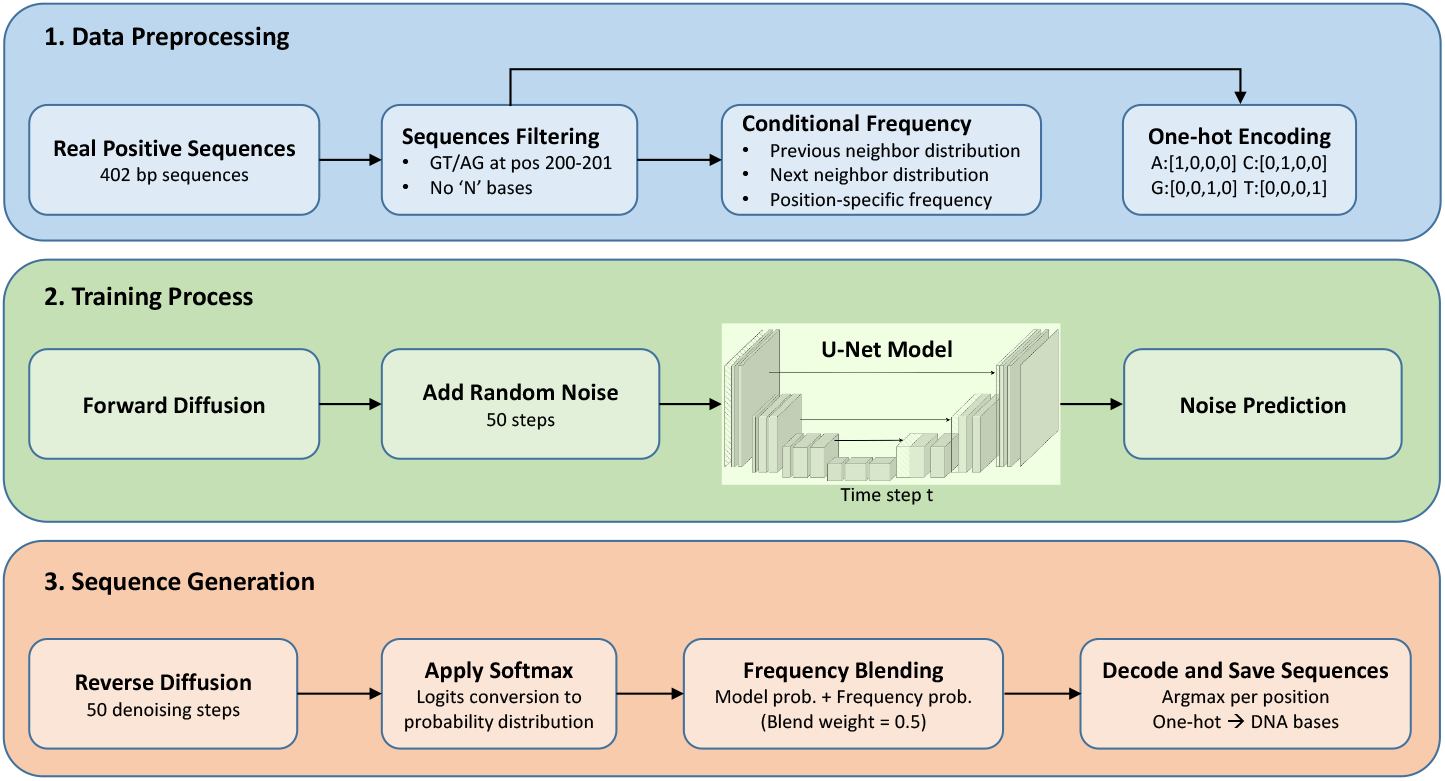
Three-stage model pipeline showing: (1) data preprocessing of real positive sequences, (2) training with forward diffusion through a U-Net architecture for noise prediction, and (3) sequence generation via reverse diffusion, frequency blending, and decoding to produce synthetic DNA sequences.

## II. Experimental Setup

### A. Data Preparation

We used the datasets from DRANetSplicer [10], which provides annotated donor and acceptor splice site sequences for *Arabidopsis thaliana* and *Homo sapiens*. Each species dataset includes four predefined sets: positive donor, positive acceptor, negative donor, and negative acceptor sequences. Positive sets contain real splice sites; negative sets contain decoy sites where a `GT`(donor) or `AG`(acceptor) motif occurs without an annotated junction.

Sequences are 402 nucleotides long with the candidate splice site centered on positions 201 and 202, providing 200 nucleotides of upstream and downstream context. All examples are aligned to center the candidate motifs.

For training of the diffusion model, we randomly selected 50,000 positive sequences per set for *Arabidopsis thaliana* and 100,000 for *Homo sapiens*. We also randomly selected equal numbers from the negative sets, which were not used for diffusion model training but later served in indirect evaluation experiments.

### B. Conditional Frequency Analysis

To guide the diffusion model towards biologically plausible sequences, we computed conditional nucleotide frequency distributions from real splice site sequences. For a DNA sequence 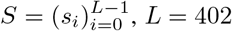, with *s_i_ ∈* {*A, C, G, T*}, we defined:

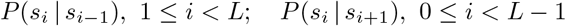

for previous and next-neighbor conditioning, respectively. The conditional probability is:

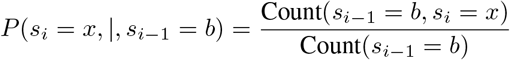

where *b, x ∈* {*A, C, G, T*}. Next-neighbor conditioning uses an analogous formula.

We generated two tables: the Previous Neighbor Table *P*_prev_(*i* − 1*, b, x*) and the Next Neighbor Table *P*_next_(*i*+1*, b, x*), where *i* indexes the position of the base to be predicted, *b* is the conditioning base and *x* the predicted nucleotide. These context-aware priors bias the diffusion model to preserve realistic dependencies between adjacent nucleotides.

### C. Diffusion Model Architecture

We employed a U-Net-based architecture [11] adapted for 1D DNA sequences as the denoising network in our diffusion framework. The model operates on one-hot encoded DNA sequences of shape (64, 402, 4), where 64 corresponds to the batch size, 402 is the sequence length, and 4 corresponds to the nucleotide channels (`A`, `C`, `G`, `T`).

The U-Net architecture consists of:

- An initial double convolution block applying two sequential 1D convolutional layers, each followed by batch normalization and rectified linear unit (ReLU) activation.
- Three downsampling blocks, each with 1D max pooling followed by a double convolution, progressively reducing sequence length while increasing feature channels.
- A bottleneck layer where a learned time-step embedding (from diffusion timestep *t*) is added to condition the model on the current noise level.
- Three upsampling blocks, each with 1D transposed convolution (stride 2) for upsampling, followed by a double convolution, with skip connections from corresponding downsampling layers.
- A final 1D convolutional layer projecting feature maps back to four nucleotide channels.

All convolutional layers use kernel size 3 with padding to maintain spatial dimensions. This architecture captures both local nucleotide motifs and long-range dependencies essential for realistic DNA sequence generation.

### D. Model Training and Synthetic Sequence Generation

The 1D U-Net diffusion model was trained to predict the noise added to one-hot encoded sequences during the forward noise process. At each step, a clean sequence *x*_0_ was perturbed to a noisy version *x_t_* according to a predefined linear noise schedule over *T* = 50 timesteps. The model received *x_t_* and the corresponding timestep *t* as inputs and was trained to minimize the mean squared error (MSE) between the predicted and true noise.

Training was conducted for 50 epochs using the Adam optimizer [12] with a learning rate of 10^−4^ and batch size of 64. Independent models were trained for donor and acceptor sites of each species. After completing the reverse diffusion process (at timestep *t* = 0), a frequency blending step refined the nucleotide probabilities using empirical conditional frequencies.

Let *p*_model_(*i*) *∈* ℝ^4^ be the model-predicted probability vector over nucleotides {*A, C, G, T*} at position *i* obtained after applying the softmax function, and *p*_freq_(*i*) *∈* ℝ^4^ be the empirical nucleotide frequency vector estimated from neighboring bases using conditional frequency tables. The final blended probability is:

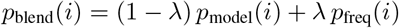

with blending weight *λ* = 0.5, chosen to balance the influence of model predictions and empirical frequencies and prevent either from dominating the generation process. The nucleotide at each position *i* is then selected as:

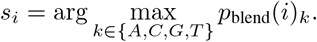

Using this procedure, we generated 60,000 synthetic sequences for each set: *Arabidopsis thaliana* donor, *Arabidopsis thaliana* acceptor, *Homo sapiens* donor, and *Homo sapiens* acceptor sequences.

## III. Results

We conducted both direct and indirect evaluations of our synthetic DNA sequences. Direct evaluation compared sequence characteristics between synthetic and real genomic data using sequence logos, `GC`content, and nucleotide conservation scores, allowing us to assess how accurately our models capture essential biological patterns and motifs.

Our direct evaluation reveals critical differences between generation methods. Sequence logos (Figures 2–3) show that while sequences generated without frequency blending (No-Blend sequences) contain the canonical `GT`/`AG`splice site dinucleotides, they fail to capture essential surrounding motifs that real genomic data displays. Frequency blending successfully reproduces species-specific patterns in these regulatory regions.

**Fig. 2.**
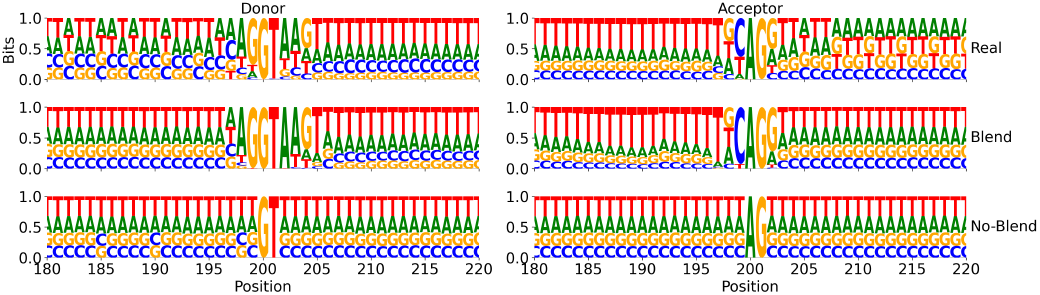
Arabidopsis thaliana sequence logos.

**Fig. 3.**
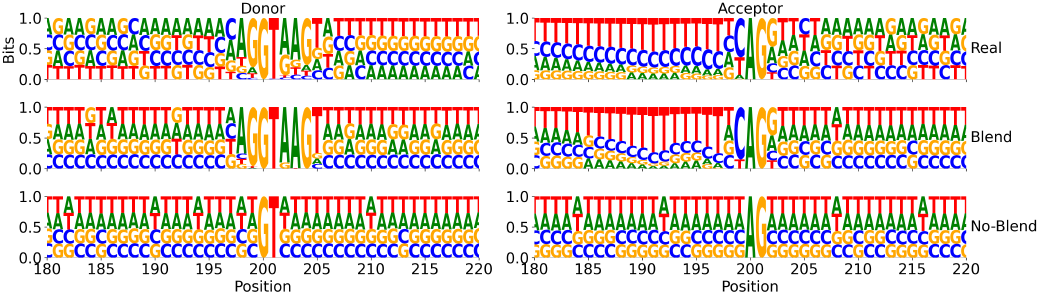
Homo sapiens sequence logos.

`GC`content analysis (Figure 4) shows that Blend sequences closely match real distributions (deviation within 0.4% in *Arabidopsis thaliana* and 0.7% in *Homo sapiens*), whereas No-Blend sequences show larger deviations: 2.5–2.7% in *Arabidopsis thaliana* and 1.5% in *Homo sapiens*. Conservation score plots (Figure 5) further confirm that Blend sequences accurately reproduce extended conservation patterns in splice site flanking regions, particularly the polypyrimidine tract preceding acceptor sites. These findings confirm that our frequency-blended diffusion models generate synthetic DNA sequences with realistic splice sites while maintaining model transferability.

**Fig. 4.**
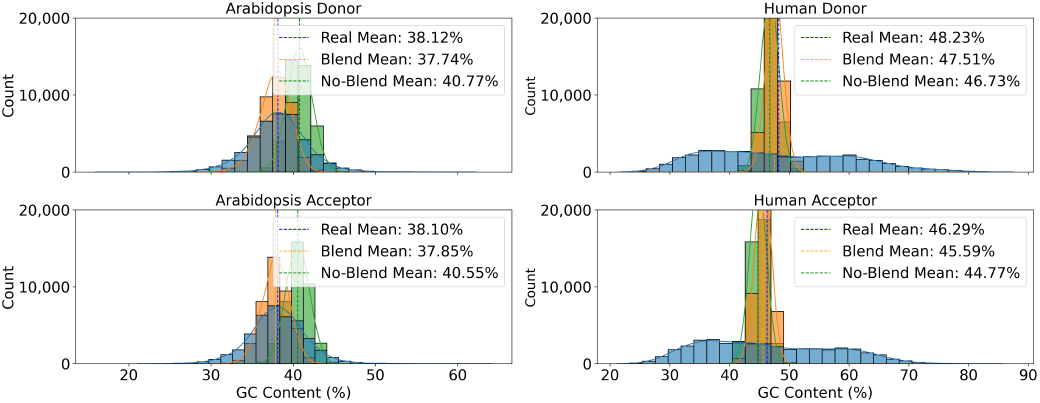
GC content.

**Fig. 5.**
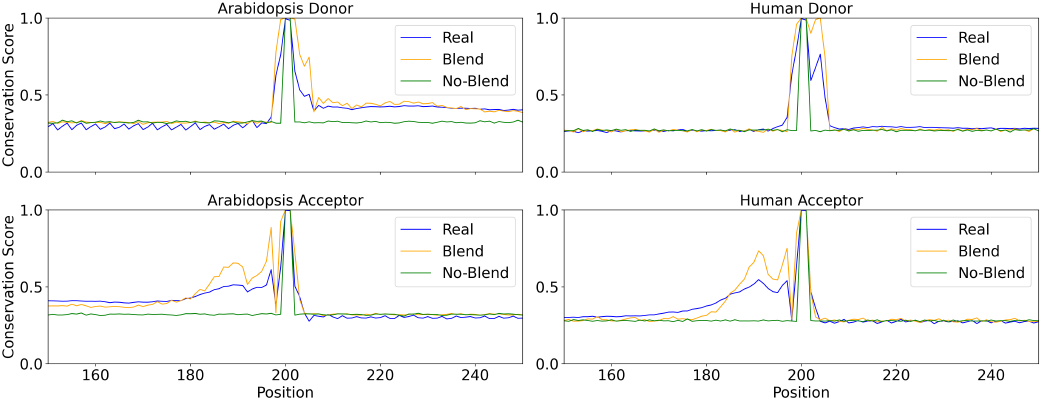
Nucleotide conservation.

For indirect evaluation, we tested the functional realism of our synthetic sequences using a model transferability framework. Two scenarios were considered: Train-Real-Test-Real (TRTR), where models were trained and tested on real data to establish baseline performance, and Train-Real-Test-Synthetic (TRTS), where models trained on real data were tested on synthetic sequences to evaluate generalization.

In TRTR, models were trained using balanced datasets consisting of 40,000 positive sequences (real splice sites) and 40,000 negative sequences (decoy sites) for training, with 10,000 positives and 10,000 negatives each for validation and testing. The best-performing model was then saved and used in TRTS, where synthetic sequences served as the positive test set and negative test sequences were taken from the remaining real data not used during training or validation.

We evaluated model effectiveness using accuracy 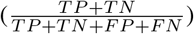, sensitivity 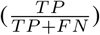, and F1score 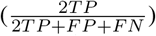 across four established splice site prediction architectures: SpliceRover [3], SpliceFinder [13], DeepSplicer [14], and Spliceator [15]. Accuracy provides an overall measure of prediction correctness, sensitivity measures the ability of a model to correctly identify true splice sites (true positives), and F1-score summarizes the trade-off between precision 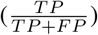 and recall (which is equivalent to sensitivity).

Results showed that TRTR performance was consistently strong (accuracy *>*0.93). However, in TRTS, models failed when frequency blending was not applied, with accuracy near random (0.52–0.54) and F1-scores below 0.21. Our frequency blending approach substantially improved results, with donor site performance recovering to real-data levels (accuracy *>*0.96) and acceptor sites also showing marked improvement. Full results for *Arabidopsis thaliana* and *Homo sapiens* are presented in Tables I and II.

These results confirm that our frequency-blended diffusion framework generates synthetic splice site sequences that closely match real genomic patterns in both structure and function. By combining learned representations with empirical frequency guidance, the model produces biologically realistic sequences suitable for training, benchmarking, and crossspecies applications.

**TABLE I.**
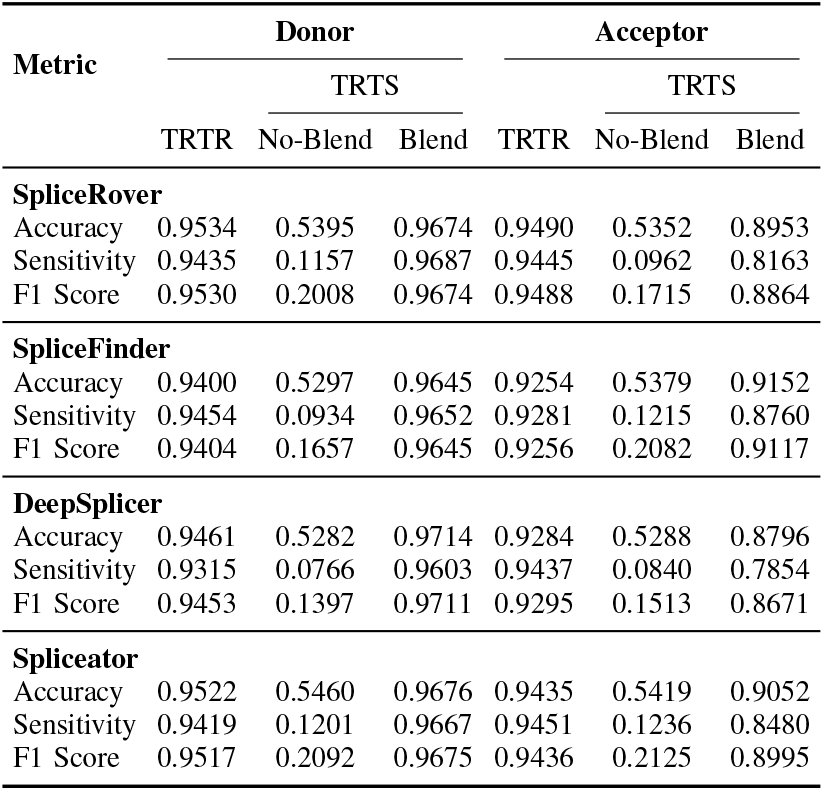
Results for *Arabidopsis thaliana*.

**TABLE II.** Results for *Homo sapiens*.

| Metric | Donor |  |  | Acceptor |  |  |
| --- | --- | --- | --- | --- | --- | --- |
|  | TRTS |  |  | TRTS |  |  |
|  | TRTR | No-Blend | Blend | TRTR | No-Blend | Blend |
| **SpliceRover** |  |  |  |  |  |  |
| Accuracy | 0.9684 | 0.5349 | 0.9710 | 0.9569 | 0.5177 | 0.8040 |
| Sensitivity | 0.9743 | 0.1111 | 0.9815 | 0.9451 | 0.0803 | 0.6527 |
| F1 Score | 0.9685 | 0.1928 | 0.9713 | 0.9564 | 0.1427 | 0.7690 |
| **SpliceFinder** |  |  |  |  |  |  |
| Accuracy | 0.9543 | 0.5332 | 0.9692 | 0.9217 | 0.5117 | 0.8457 |
| Sensitivity | 0.9571 | 0.1206 | 0.9892 | 0.8999 | 0.0906 | 0.7565 |
| F1 Score | 0.9544 | 0.2053 | 0.9698 | 0.9200 | 0.1565 | 0.8305 |
| **DeepSplicer** |  |  |  |  |  |  |
| Accuracy | 0.9570 | 0.5219 | 0.9613 | 0.9425 | 0.5232 | 0.7930 |
| Sensitivity | 0.9540 | 0.0992 | 0.9782 | 0.9514 | 0.1042 | 0.6470 |
| F1 Score | 0.9569 | 0.1718 | 0.9619 | 0.9430 | 0.1793 | 0.7576 |
| **Spliceator** |  |  |  |  |  |  |
| Accuracy | 0.9635 | 0.5248 | 0.9721 | 0.9492 | 0.5176 | 0.8071 |
| Sensitivity | 0.9771 | 0.0872 | 0.9812 | 0.9521 | 0.0766 | 0.6522 |
| F1 Score | 0.9639 | 0.1550 | 0.9724 | 0.9493 | 0.1370 | 0.7717 |

## IV. Conclusions and Future Work

In this study, we introduced a frequency-blended diffusion model for generating synthetic DNA sequences containing biologically realistic splice sites. By integrating deep generative modeling with conditional nucleotide frequency priors, our approach effectively captured both local sequence motifs and long-range dependencies. Through comprehensive evaluations, we demonstrated that our synthetic sequences closely matched real genomic patterns in composition, conservation, and functional predictability.

Future work will explore applying frequency blending to other generative models such as generative adversarial networks (GANs), variational autoencoders (VAEs), and flow-matching methods. We also aim to extend the framework to additional biological tasks beyond splice sites, support multispecies and multitask generation, and integrate functional constraints like predicted splicing or regulatory effects to improve biological relevance.

## Appendix for Frequency-Blended Diffusion for Splice Site Sequence Generation

### Comparison to the original U-Net

Our model design is inspired by the classical U-Net architecture introduced for biomedical image segmentation [11]. While the original U-Net uses a 2D encoder-decoder structure with Conv2D and MaxPool2D layers for pixel-wise segmentation of images, our model adapts this concept to 1D sequential data to suit DNA sequences. Specifically, we replace 2D operations with Conv1D, MaxPool1D, and UpConv1D layers to process one-hot encoded DNA sequences of length 402 with four nucleotide channels. We maintain the core architectural principles of U-Net: an encoder-decoder structure with skip connections and progressive channel expansion and reduction. However, our model introduces two key innovations: (1) the incorporation of a learned diffusion timestep embedding, which conditions the bottleneck features during denoising, and (2) the integration of a frequency blending step, where the model output is combined with empirical conditional nucleotide frequency priors to guide biologically realistic sequence generation. Additionally, unlike the original U-Net, which uses unpadded convolutions and requires cropping, our model employs padding to maintain consistent feature map sizes. Thus, our model can be considered a 1D diffusion-conditioned variant of U-Net specifically adapted for synthetic DNA sequence generation.

### Comparison between our work and DNA-Diffusion

Both our study and the DNA-Diffusion [8] approach leverage diffusion probabilistic models with U-Net-based denoising architectures for synthetic DNA sequence generation. However, there are key architectural and methodological differences:

#### A. Sequence type and length

DNA-Diffusion focuses on generating 200 bp regulatory elements (derived from DNase I Hypersensitive Site (DHS) peak summits), while our model generates full-length 402 bp splice site-centered sequences.

#### B. Model conditioning

DNA-Diffusion introduces two conditioning signals into U-Net: a continuous time-step embedding and a discrete cell-type label embedding to enable multi-cell type generation. Our model only conditions on the diffusion timestep using a learned time-step embedding, as the splice site sequences are species- and site-specific and do not require cell-type conditioning.

#### C. U-Net architecture

The DNA-Diffusion model uses a larger and more complex U-Net with:

- 200 input channels,
- downsampling stages with ResNet blocks and linear attention layers,
- upsampling stages with mirroring architecture and additional ResNet blocks before the output layer.

Our model uses a simpler and lightweight 1D U-Net optimized for nucleotide sequences:

- Input: one-hot encoded sequences of shape (64, 402, 4),
- 3 downsampling blocks (max pooling + double 1D convolution),
- a bottleneck block with addition of time embedding,
- 3 upsampling blocks (transposed convolution + double convolution),
- final 1D convolution to predict denoised sequence logits.

#### D. Additional guidance strategy

A key innovation of our approach is the **conditional frequency blending** step applied after the reverse diffusion process. Nucleotide probabilities predicted by the diffusion model are blended with position-specific conditional nucleotide frequencies estimated from real sequence data to enforce local sequence realism. DNA-Diffusion does not implement such external statistical guidance.

#### E. Model objective and application

DNA-Diffusion aims to design regulatory elements to modulate chromatin accessibility and gene expression across multiple cell types. Our model specifically targets the generation of biologically realistic splice site regions for *Arabidopsis thaliana* and *Homo sapiens*, to augment datasets for splice site prediction.

### Conditional Frequency Table Computation

To guide the diffusion model toward biologically realistic sequences, we computed conditional nucleotide frequency distributions from real splice site sequence alignments. These distributions represent the probability of observing a nucleotide at a given position conditioned on the nucleotide in the adjacent position, either upstream (previous neighbor) or downstream (next neighbor).

For each sequence *S* = (*s*_0_*, s*_1_*, …, s*_*L*−1_) of length *L* = 402, with *s_i_ ∈* {*A, C, G, T*}, we define:

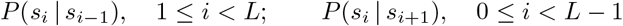

for previous- and next-neighbor conditioning, respectively. The conditional probability is estimated as:

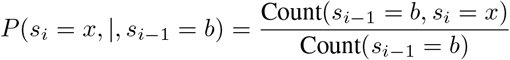

where *b, x ∈* {*A, C, G, T*}. Next-neighbor probabilities are computed analogously. These distributions are stored as:

- Previous neighbor table: *P*_prev_(*i* − 1*, b, x*)
- Next neighbor table: *P*_next_(*i* + 1*, b, x*)

where *i* denotes the position of the current base, *b* is the conditioning base, and *x* is the predicted nucleotide.

Table A1 illustrates an example excerpt of a previous-neighbor conditional frequency table computed from real donor site sequences (*Arabidopsis* dataset, positions 198–202).

**TABLE A1.** Example of previous-neighbor conditional frequency table for donor sequences (*Arabidopsis*).

| Position | Previous base ($b$) | Current base ($x$) | Probability $P_{\text{prev}}(i - 1, b, x)$ |
| --- | --- | --- | --- |
| 200 | G | T | 1.00 |
| 199 | A | G | 0.85 |
| 199 | A | C | 0.10 |
| 199 | A | A | 0.05 |
| 201 | T | C | 0.60 |
| 201 | T | A | 0.40 |

This position-specific conditional frequency information is subsequently used during generation to blend model predictions with empirically observed nucleotide patterns, improving biological plausibility of the synthetic sequences.

### Blending Model Output with Conditional Frequencies

At the end of the reverse diffusion process, the model outputs a raw prediction tensor representing nucleotide scores (logits) for each position. To interpret these as probabilities, a softmax function is applied across the four nucleotide channels at each position:

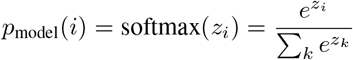

where *z_i_* is the vector of predicted scores at position *i* and *∈* {*k A, C, G, T*}.

These softmax-normalized probabilities represent the confidence of the model over possible nucleotides at each position. However, model outputs alone may still exhibit biologically implausible patterns. To mitigate this, we blend the model predictions with empirical conditional nucleotide frequencies derived from real splice site datasets. These conditional frequencies act as a guide reflecting species-specific sequence constraints.

The final nucleotide probability vector at position *i* is calculated as:

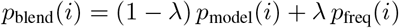

where *p*_freq_(*i*) is the conditional frequency vector, and *λ* = 0.5 is the blending weight.

Table A2 shows an illustrative example at position *i*. Suppose the diffusion model strongly predicts `G`as the most likely nucleotide, but biological data suggests both `C`and `G`are common. Blending incorporates both signals. In this example:

- The model predicts `G`with high confidence (0.60), while real data suggests `C`and `G`are both common (0.30 and 0.40).
- Blending reduces overconfidence in `G`and slightly increases probabilities for `C`and `A`.
- The final nucleotide is selected as the one with the highest blended probability (here, `G`at 0.50).

**TABLE A2.** Example of nucleotide probability blending at position *i*.

| Nucleotide | Model Prediction | Frequency Prior | Blended Probability |
| --- | --- | --- | --- |
| A | 0.10 | 0.20 | 0.15 |
| C | 0.15 | 0.30 | 0.225 |
| G | 0.60 | 0.40 | 0.50 |
| T | 0.15 | 0.10 | 0.125 |

This blending strategy provides a soft correction mechanism, balancing learned model outputs with biologically grounded priors to generate sequences that better reflect real species-specific patterns.

Table A3 summarizes the main hyperparameters used during training and generation for our frequency-blended diffusion model.

**TABLE A3.** Hyperparameters for model training and sequence generation.

| Parameter | Value |
| --- | --- |
| Model architecture | 1D U-Net (3 downsampling + 3 upsampling blocks) |
| Base channels | 32 |
| Diffusion timesteps | 50 |
| Optimizer | Adam |
| Learning rate | $1 \times 10^{-4}$ |
| Batch size | 64 |
| Epochs | 50 |
| Blending weight $\lambda$ | 0.5 |

All models were trained independently for donor and acceptor splice sites for each species using the same hyperparameters. Blending was applied only at generation time to softly correct model outputs using empirical conditional nucleotide frequencies.

### Indirect Evaluation Proxy Models

We used four established convolutional neural network architectures as proxy models to evaluate the realism of our synthetic splice site sequences in the indirect evaluation experiments. All models take as input one-hot encoded DNA sequences of shape (64, 402, 4) and output a binary classification prediction for splice site detection.

a. *SpliceRover [3].:* SpliceRover consists of five sequential 1D convolutional blocks with increasing channel depth: Conv1D layers (70, 100, 100, 200, 250 filters), kernel sizes of 9 or 7, ReLU activations, dropout (0.2), and max pooling layers in the last three blocks. A fully connected layer (512 neurons) followed by a softmax output layer performs classification.
b. *SpliceFinder [13].:* SpliceFinder uses a single Conv1D layer with 50 filters and kernel size of 9, followed by ReLU activation. The output is flattened and passed to two dense layers (100 and 2 neurons), with dropout (0.3) after the first dense layer.
c. *DeepSplicer [14].:* DeepSplicer applies three stacked Conv1D layers with 50 filters and kernel size of 9 in each layer. Each convolutional block uses ReLU activation. The flattened output is passed to two dense layers (100 and 2 neurons), with dropout (0.3) after the first dense layer.
d. *Spliceator [15].:* Spliceator is a compact CNN with three Conv1D blocks: 16 filters (kernel size 7), 32 filters (kernel size 6), and 64 filters (kernel size 6), each followed by ReLU activation, max pooling, and dropout (0.2). The final dense layers consist of 100 neurons and 2 output neurons.

All models were trained using the Adam optimizer (learning rate 0.001) and cross-entropy loss. Each architecture was evaluated in both Train-Real-Test-Real (TRTR) and Train-Real-Test-Synthetic (TRTS) scenarios to assess the generalization of models trained on real data when applied to synthetic sequences.

Table A4 provides a summarizing comparison of the architectural characteristics of these proxy models.

**TABLE A4.** Detailed layer-by-layer architecture of the proxy models used for indirect evaluation.

| Layer | SpliceRover | SpliceFinder | DeepSplicer | Spliceator |
| --- | --- | --- | --- | --- |
| 1 | Conv1D (70, 9) | Conv1D (50, 9) | Conv1D (50, 9) | Conv1D (16, 7) |
| 2 | Dropout (0.2) | Flatten | Conv1D (50, 9) | MaxPool (2) |
| 3 | Conv1D (100, 7) | Dense (100) | Conv1D (50, 9) | Dropout (0.2) |
| 4 | Dropout (0.2) | Dropout (0.3) | Flatten | Conv1D (32, 6) |
| 5 | Conv1D (100, 7) | Dense (2) | Dense (100) | MaxPool (2) |
| 6 | MaxPool (3) | - | Dropout (0.3) | Dropout (0.2) |
| 7 | Dropout (0.2) | - | Dense (2) | Conv1D (64, 6) |
| 8 | Conv1D (200, 7) | - | - | MaxPool (2) |
| 9 | MaxPool (4) | - | - | Dropout (0.2) |
| 10 | Dropout (0.2) | - | - | Flatten |
| 11 | Conv1D (250, 7) | - | - | Dense (100) |
| 12 | MaxPool (4) | - | - | Dense (2) |
| 13 | Dropout (0.2) | - | - | - |
| 14 | Flatten | - | - | - |
| 15 | Dense (512) | - | - | - |
| 16 | Dropout (0.2) | - | - | - |
| 17 | Dense (2) | - | - | - |

All proxy models were evaluated using the same experimental pipeline to ensure comparability across architectures. Table A5 summarizes the key hyperparameters used.

**TABLE A5.**
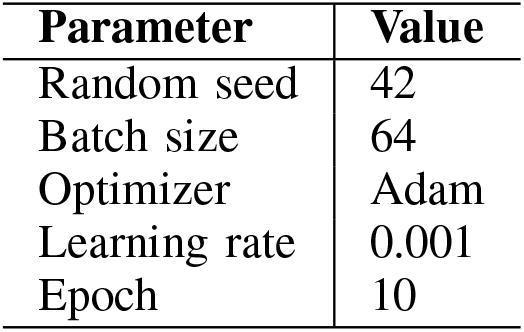
Hyperparameters for proxy model evaluation experiments.

## References

[1] G. Martín, Y. Márquez, F. Mantica, P. Duque, and M. Irimia, “Alternative splicing landscapes in Arabidopsis thaliana across tissues and stress conditions highlight major functional differences with animals,” Genome Biology, vol. 22, no. 1, p. 35, 2021.

[2] Y. Xing and C. Lee, “Alternative splicing and RNA selection pressure– evolutionary consequences for eukaryotic genomes,” Nature Reviews Genetics, vol. 7, no. 7, pp. 499–509, 2006.

[3] J. Zuallaert, F. Godin, M. Kim, A. Soete, Y. Saeys, and W. De Neve, “SpliceRover: interpretable convolutional neural networks for improved splice site prediction,” Bioinformatics, vol. 34, no. 24, pp. 4180–4188, 2018.

[4] M. Pop, T. K. Attwood, J. A. Blake, P. E. Bourne, A. Conesa, T. Gaasterland, L. Hunter, C. Kingsford, O. Kohlbacher, T. Lengauer, S. Markel, Y. Moreau, W. S. Noble, C. Orengo, B. F. F. Ouellette, L. Parida, N. Przulj, T. M. Przytycka, S. Ranganathan, R. Schwartz, A. Valencia, and T. Warnow, “Biological databases in the age of generative artificial intelligence,” Bioinformatics Advances, vol. 5, no. 1, p. vbaf044, 03 2025.

[5] W. Ge, M. Meier, C. Roth, and J. Söding, “Bayesian Markov models improve the prediction of binding motifs beyond first order,” NAR Genomics and Bioinformatics, vol. 3, no. 2, p. lqab026, 04 2021.

[6] M. Siebert and J. Söding, “Bayesian Markov models consistently out-perform PWMs at predicting motifs in nucleotide sequences,” Nucleic acids research, vol. 44, no. 13, pp. 6055–6069, 2016.

[7] F. Calvanese, G. Peinetti, P. Pavlinova, P. Nghe, and M. Weigt, “Integrating experimental feedback improves generative models for biological sequences,” arXiv preprint arXiv:2504.01593, 2025.

[8] L. F. DaSilva, S. Senan, Z. M. Patel, A. J. Reddy, S. Gabbita, Z. Nussbaum, C. M. V. Córdova, A. Wenteler, N. Weber, T. M. Tunjic et al., “DNA-Diffusion: Leveraging Generative Models for Controlling Chromatin Accessibility and Gene Expression via Synthetic Regulatory Elements,” bioRxiv, 2024.

[9] Z. Li, Y. Ni, T. Huygelen, A. Das, G. Xia, G.-B. Stan, and Y. Zhao, “Latent Diffusion Model for DNA Sequence Generation,” in NeurIPS 2023 AI for Science Workshop, 2023.

[10] X. Liu, H. Zhang, Y. Zeng, X. Zhu, L. Zhu, and J. Fu, “DRANetSplicer: A Splice Site Prediction Model Based on Deep Residual Attention Networks,” Genes, vol. 15, no. 4, p. 404, 2024.

[11] O. Ronneberger, P. Fischer, and T. Brox, “U-Net: Convolutional Networks for Biomedical Image Segmentation,” in Medical Image Computing and Computer-Assisted Intervention – MICCAI 2015. Springer, 2015, pp. 234–241.

[12] D. P. Kingma and J. Ba, “Adam: A Method for Stochastic Optimization,” arXiv preprint arXiv:1412.6980, 2014.

[13] R. Wang, Z. Wang, J. Wang, and S. Li, “SpliceFinder: ab initio prediction of splice sites using convolutional neural network,” BMC bioinformatics, vol. 20, pp. 1–13, 2019.

[14] V. Akpokiro, O. Oluwadare, and J. Kalita, “DeepSplicer: An Improved Method of Splice Sites Prediction using Deep Learning,” in 2021 20th IEEE international conference on machine learning and applications (ICMLA). IEEE, 2021, pp. 606–609.

[15] N. Scalzitti, A. Kress, R. Orhand, T. Weber, L. Moulinier, A. Jeannin-Girardon, P. Collet, O. Poch, and J. D. Thompson, “Spliceator: multi-species splice site prediction using convolutional neural networks,” BMC bioinformatics, vol. 22, pp. 1–26, 2021.

